# VANGL2 shapes the mouse heart tube from adjacent epithelia and without planar polarity

**DOI:** 10.1101/2025.09.05.674213

**Authors:** Paul Palmquist-Gomes, Gaëlle Letort, Ayushi U. Hegde, José María Pérez-Pomares, Sigolène M. Meilhac

## Abstract

Disruption of the core Planar Cell Polarity component VANGL2 in mice is associated with congenital heart defects and impaired morphogenesis of the embryonic heart tube. However, the underlying mechanisms have remained unclear. Here we quantified in 3D the heart geometry and adjacent tissue architecture in a series of mutants to reveal a dual role of *Vangl2* in shaping the heart tube. Together with cell labelling in the chick, we show that VANGL2 in multicellular junctions promotes second heart field cell rearrangements and thus the elongation of the arterial pole. In addition, apically localised VANGL2 and its downstream actin-binding effector SHROOM3 control the bilateral symmetry of the splanchnic mesoderm caudal to the venous pole. Disorganisation of this midline anchor is associated with a rotation of the heart tube and abnormal left ventricle position. Our work overall uncovers novel specific epithelial roles of VANGL2 unrelated to planar polarity during heart morphogenesis.

## Introduction

Epithelial tissues are crucial in morphogenesis. They serve as platforms of organised cells with specific junctions and polarity, key to orient tissue deformation^1^. Cell behaviour is collectively coordinated in epithelia, for example driving convergent extension or specific folding of the cell sheet. During convergent extension, cell rearrangements such as cell intercalation, division or shape changes are coordinated to elongate the tissue specifically along one axis, at the expense of the perpendicular axis. An example of one such mechanism is the directional contraction of cell-cell interfaces leading to the formation of multicellular rosette structures, which resolve in new cell-cell interfaces in a perpendicular direction^2^. For tissue folding, epithelial sheets require mechanical asymmetries, which can be generated locally by apical cell constriction^3^. Although these processes have been extensively studied during fly development, using easy-access genetics and live-imaging, assessing their conservation during mammalian organogenesis remains challenging.

The planar cell polarity (PCP) pathway is an important regulator of cell behaviour coordination across epithelia. Canonical PCP relies on opposing protein complexes in abutting cell-cell interfaces, involving the transmembrane proteins FRIZZLED, CELSR and VAN GOGH (VANGL), and cytosolic proteins DISHEVELLED, ANKRD6 and PRICKLE, which polarise cells in the epithelium plane^4–6^. In mammals, the multisystemic phenotype of the spontaneous loop-tail (Lp) mutant indicates that *Vangl2* is required for morphogenesis of several organs, including the cochlea, neural tube, lung^7–10^, and for convergent extension of the body axis^11^. During neural tube closure, VANGL2 has been shown to act upstream of the actin-binding protein SHROOM3 to control apical constriction and folding of the neural plate^12–14^. VANGL2 and its orthologue VANG-1 are essential for multicellular rosette resolution underlying convergent extension of the mouse neural plate and *C. elegans* ventral nerve cord, respectively^15,16^. VANGL2 can also act independently of other core PCP components to maintain smooth muscle architecture^10^. While *Vangl2* is conserved between animal species, it plays diverse tissue-specific roles.

*Vangl2* is required for heart morphogenesis. Loss of *Vangl2* in mice leads to congenital heart defects, such as double-outlet right ventricle, double-sided aortic arch and coronary vasculature defects^17,18^. *Shroom3* constitutive mutants have double-outlet right ventricle, similar to *Vangl2* mutants^19^. During development, *Vangl2* inactivation has been associated with abnormal heart looping and shortening of the outflow tract^20^. However, the cellular mechanisms underlying these defects have remained incompletely understood.

Because *Vangl2* is required for neural tube morphogenesis, it was initially suggested that defects in the heart were secondary to that in the neural tube^17^. However, *Vangl2* is expressed in cardiac progenitors, as well as in the myocardium^20,21^. Conditional inactivation has shown that *Vangl2* is required for outflow tract elongation in the heart field labelled by *Isl1^Cre^*, and not in the myocardial or neural crest lineages^20^. Derived from the splanchnic lateral plate mesoderm, the second heart field is an epithelium that covers the dorsal pericardial wall behind the embryonic heart and contains cardiac precursors, which feed the heart tube while it grows^22,23^. The heart field is continuous with the arterial and venous poles of the heart tube dorsally, while ventrally these poles are connected to the pericardium or the sinus venosus. Elongation of the heart tube takes place while the arterial and venous poles are fixed in place, so that it deforms by buckling during the process of heart looping at E8.5^24^. Heart looping is modulated by the breakdown of the dorsal mesocardium that initially anchors the heart tube at the midline^24^. Additionally, left-right patterning controls the orientation of buckling, leading to a rightward helical shape of the looped heart tube^25^. During looping, the right ventricle is repositioned from cranial to right. In general, looping is important to align cardiac chambers for the establishment of the double blood circulation. The heart tube continues to elongate after heart looping, until E10.5^26,27^, and is fed by other fields of precursors^28–30^. It is still unclear how *Vangl2* regulates heart tube morphogenesis and in which source of progenitors.

The second heart field is polarised along an apico-basal axis, facing the pericardial cavity and the endoderm, respectively^31^. Expression of PCP genes *Vangl2* and *Wnt5a* has been detected in the second heart field^20,32^. However, a polarised localisation of VANGL2 in the plane of the dorsal pericardial wall epithelium — which would be expected if the core PCP pathway is active — has not yet been investigated. The shorter outflow tract observed in *Vangl2* and *Prickle1* mutants^20,33^ suggests that the PCP pathway in the second heart field normally controls heart tube elongation. In chick and mouse embryos, results of cell tracing experiments are compatible with convergent extension of second heart field progenitors contributing to outflow tract elongation^34–36^. Whether the PCP pathway controls convergent extension in the second heart field has not yet been demonstrated, and the associated cell behaviour has not been characterised.

Here, we investigate the role of *Vangl2* in mouse heart tube morphogenesis. Using 3D reconstructions of the heart tube at sequential stages, we quantify defects in outflow tract length and ventricle position. Comparison of constitutive and conditional mutants reveals that *Vangl2* is required in two distinct cell populations, the *Mesp1^Cre^*-positive dorsal pericardial wall and the *Hoxb1^Cre^*-positive splanchnic mesoderm caudal to the heart tube. We tackle the challenge of imaging the subcellular localisation of VANGL2 to show that it is not planar polarised in the second heart field. By quantifying cellular architecture in mice and manipulating cell behaviour in the chick embryo, we demonstrate that *Vangl2* is rather required for multicellular rosette resolution in the second heart field, underlying outflow tract elongation. Finally, we find that inactivation of *Vangl2* or *Shroom3* disrupts the architecture of the caudal splanchnic mesoderm and impacts left ventricle position, by abnormal rotation of the heart tube around its midline anchor. Our results overall show that *Vangl2* plays dual roles in adjacent epithelia to shape the heart tube at the arterial and venous poles.

## Results

### *Vangl2* is required for heart tube shape, independently of neural tube defects

To analyse the role of *Vangl2* in the formation of the heart, we generated constitutive mutants (Fig. S1A-C) and controlled that they lack VANGL2 protein (Fig. S1E-G). As expected from observations in the spontaneous *Vangl2^Lp/Lp^* mutants^7,11,17,37^, *Vangl2^null/null^* mutants at E9.5 have neural tube and body axis elongation defects (Fig. S1A-C), as well as anomalies in heart tube shape (Fig. 1). Neural tube defects (namely, those resulting in an open neural tube) have a partial penetrance, while all mutant embryos have body axis elongation defects (Fig. S1A-C), reflecting impaired convergence-extension^11^. Heart tube defects were also detected in mutants with a normally closed neural tube (Fig. S1I), indicating that they occur independently of severe neural tube defects.

**Figure 1.**
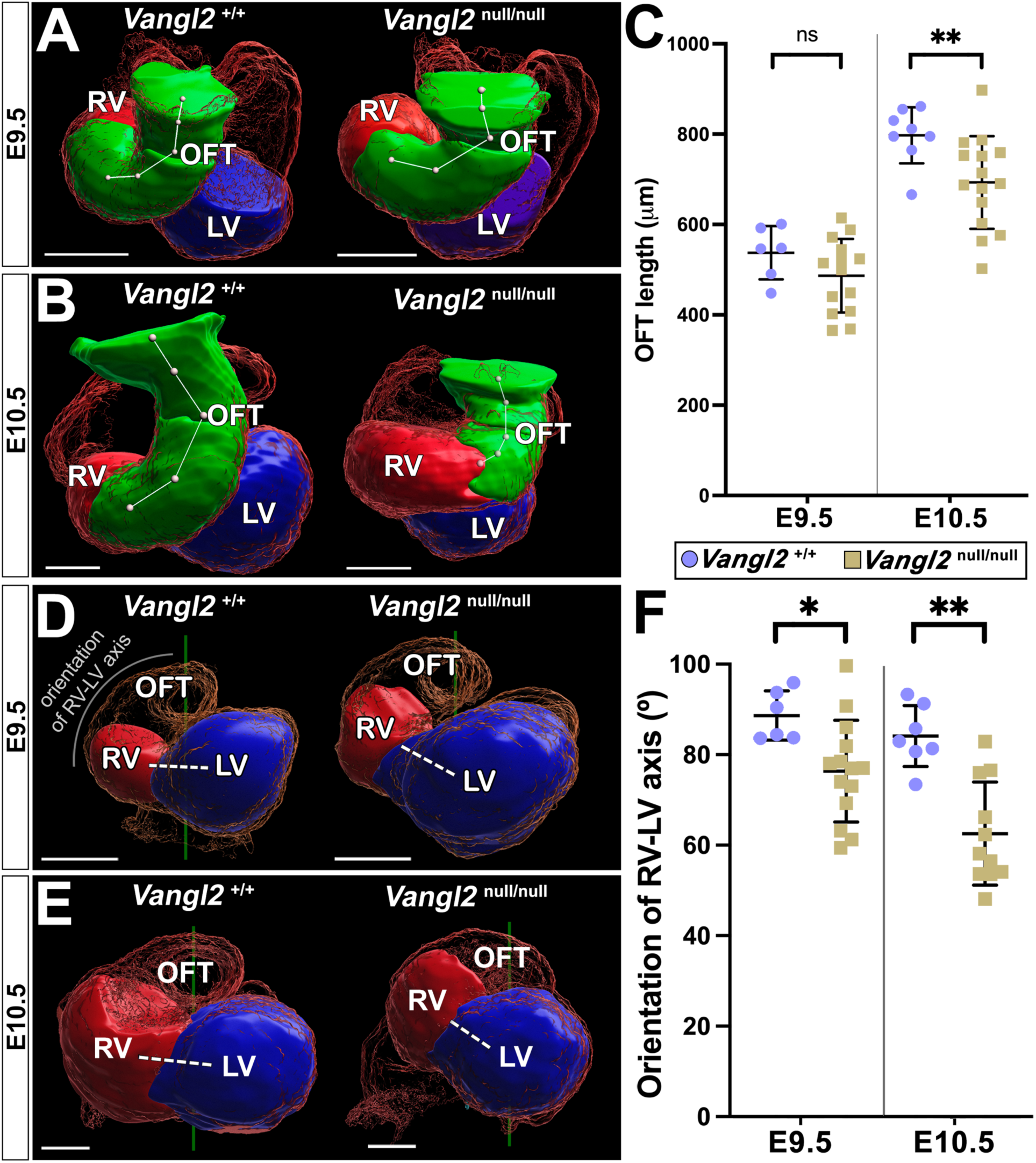
Constitutive inactivation of *Vangl2* impairs heart tube shape and outflow tract elongation. (A-B) 3D segmentation of the outflow tract (OFT, green) at E9.5 (A) and E10.5 (B), in *Vangl2* constitutive mouse mutants (*Vangl2*^null/null^) and wild-type littermates shown in a cranial view. (C) Corresponding quantification of the outflow tract length. ** p value < 0.01 (Mann Whitney t-test, n=14 and 15 *Vangl2*^null/null^, 6 and 8 *Vangl2*^+/+^ embryos at E9.5 and E10.5, respectively). (D) 3D segmentations of the right (RV, red) and left (LV, blue) ventricles at E9.5 (D) and E10.5 (E) shown in a ventral view. Hearts are aligned based on the notochord (nt, green) and dorso-ventral axis. (F) Corresponding quantification of the orientation of the RV-LV axis (dashed white line in D) relative to the notochord. * p value < 0.05, **<0.01 (Mann Whitney t-test, n=14 and 11 *Vangl2*^null/null^, 6 and 7 *Vangl2*^+/+^ embryos at E9.5 and E10.5, respectively). Means and standard deviations are shown. ns, non-significant. Scale bars : 300µm. See also Figure S1-S2, Source data.

We further quantified heart defects based on 3D imaging (Fig. 1). At E9.5, *Vangl2^null/null^* mutant hearts display a normal heart tube length and distance between the arterial and venous poles (Fig. S2A-C), indicating that the buckling mechanism, central to heart looping^24^, has proceeded normally. Yet, *Vangl2^null/null^* mutant hearts have an abnormal helical shape, which manifests as an abnormal tilted position of ventricles not parallel to the left-right axis, as well as a shorter outflow tract. Defects in the heart tube shape are more severe at E10.5 compared to E9.5 (Fig. 1C, F), showing that *Vangl2* continues to regulate heart tube morphogenesis after heart looping has ended. Our phenotyping overall shows that *Vangl2* is required for shaping the heart tube after heart looping and independently of its role in the neural tube.

### *Vangl2* in the *Mesp1-*positive mesoderm controls outflow tract elongation but not ventricle position

To understand the origin of the heart defects associated with loss of *Vangl2*, we generated *Vangl2* conditional mutants with targeted inactivation in cardiac precursors expressing *Mesp1*^38^. In keeping with the genetic tracing of *Mesp1* (Fig. S3A-G), *Mesp1*^Cre/+^;*Vangl2*^flox/null^ have no defect in the neural tube and body axis elongation (Fig. S1D, H). Similar to *Vangl2^null/null^* constitutive mutants, *Mesp1*^Cre/+^;*Vangl2*^flox/null^ conditional mutants display a significant shortening of the outflow tract (Fig. 2A-C). However, they do not show the tilted position of ventricles (Fig. 2D-F). In addition, we did not find in *Vangl2^null/null^* constitutive mutants a correlation between the outflow tract length and the angular orientation of the right ventricle-left ventricle axis (Fig. S2D). This finding demonstrate that outflow tract defects alone cannot explain the defective helical shape of the heart tube. Thus, *Vangl2* in the *Mesp1*-positive mesoderm is important for the elongation of the arterial pole but does not account for the entire cardiac phenotype observed in constitutive mutants.

**Figure 2.**
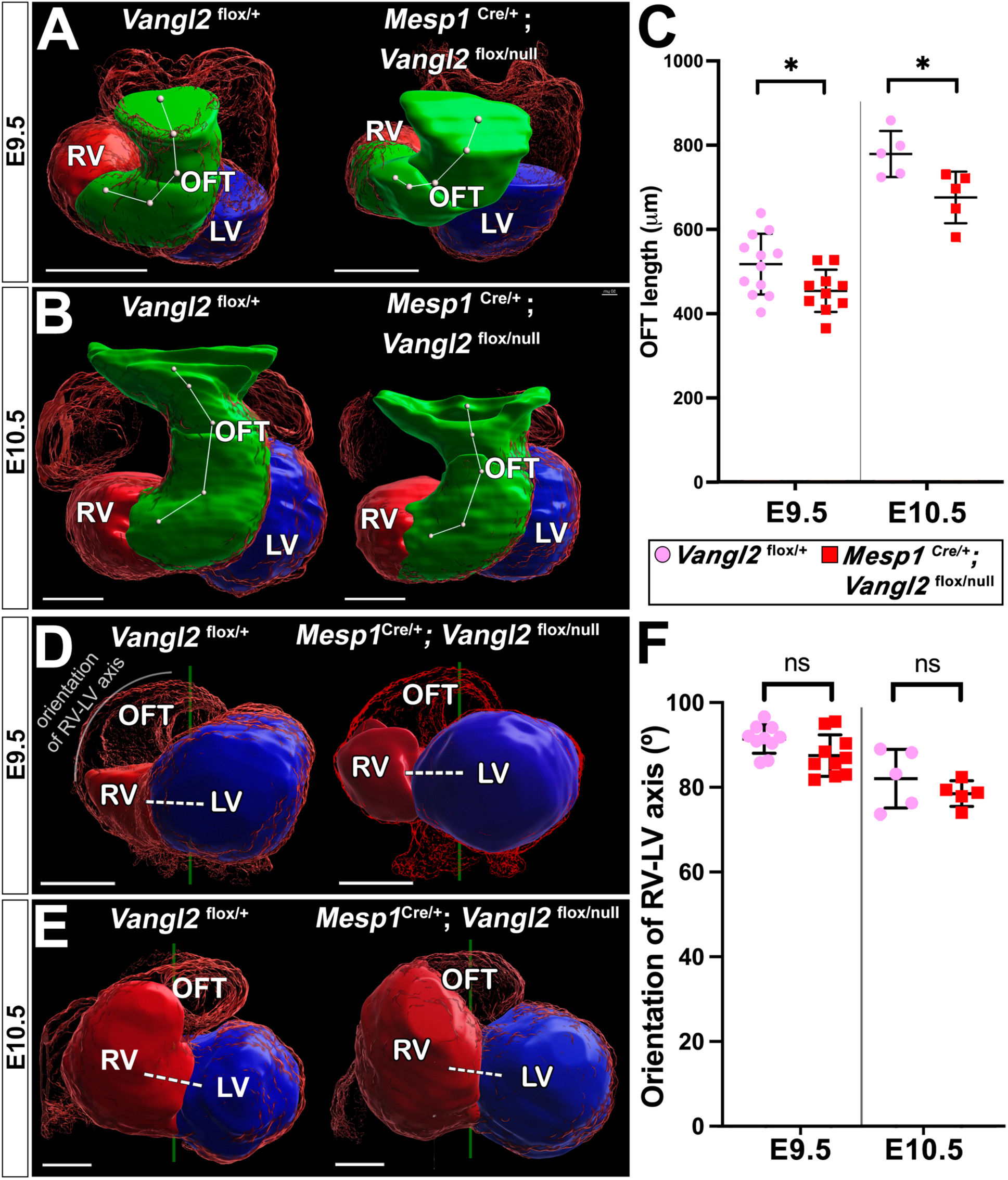
Conditional inactivation of *Vangl2* in the *Mesp1*-positive mesoderm disrupts outflow tract elongation but not ventricle position. (A-B) 3D segmentation of the outflow tract (OFT, green) at E9.5 (A) and E10.5 (B), in *Vangl2* conditional mouse mutants and control littermates shown in a cranial view. (C) Corresponding quantification of the outflow tract length. * p value < 0.05 (Mann Whitney t-test, n=10 and 5 *Mesp1*^Cre/+^;*Vangl2*^flox/null^, 11 and 5 *Vangl2*^flox/+^ embryos at E9.5 and E10.5, respectively). (D) 3D segmentations of the right (RV, red) and left (LV, blue) ventricles at E9.5 (D) and E10.5 (E) shown in a ventral view. (F) Corresponding quantification of the orientation of the RV-LV axis (dashed white line in D) relative to the notochord (green line in D). (Mann Whitney t-test, n=10 and 5 *Mesp1*^Cre/+^;*Vangl2*^flox/null^, 10 and 5 *Vangl2*^flox/+^ embryos at E9.5 and E10.5, respectively). Means and standard deviations are shown. ns, non-significant ; OFT, outflow tract. Scale bars : 300µm. See also Figure S3, Source data.

### VANGL2 is not planar polarised and accumulates in multicellular junctions of cardiac progenitors

We then analysed how *Vangl2* can control the elongation of the outflow tract. VANGL2 is involved in Planar Cell Polarity in the mouse epidermis and neural tube^6,13^. Thus, we investigated its cellular localisation in cardiac progenitors that contribute to the outflow tract, i.e. in the second heart field located within the dorsal pericardial wall at E9.5. In contrast to the neural tube, where VANGL2 levels are uniformly high throughout cell membranes (Fig. S1E), VANGL2 in cardiac progenitors is less concentrated (Fig. S1J). Unlike in the epidermis^6,39^, VANGL2 is not enriched in specific cell-cell interfaces and thus not planar polarised in cardiac progenitors (Fig. 3A-C). Instead, VANGL2 accumulates in cell vertices, specifically 4 cell- and multicellular junctions corresponding to rosettes (Fig. 3B, E-F). In the lung mesenchyme, an atypical, i.e. not planar polarised, distribution of VANGL2 is associated with an incomplete expression of core PCP pathway components^40^. We thus analysed in a published single cell transcriptomic dataset^41^ the expression levels of core PCP genes in the heart field and outflow tract (Fig. S4). In contrast to *Vangl2* and the ligand gene *Wnt5a*, *Prickle2-3* and *Celsr1-3* are not expressed, and *Prickle1* and *Wnt11* are expressed in the outflow tract but not the heart field (Fig. S4G-P). Overall, our results indicate that the core PCP pathway is not complete and thus not able to establish planar polarity in the second heart field at E9.5. This suggests that VANGL2 plays an alternative role, in multicellular junctions.

**Figure 3.**
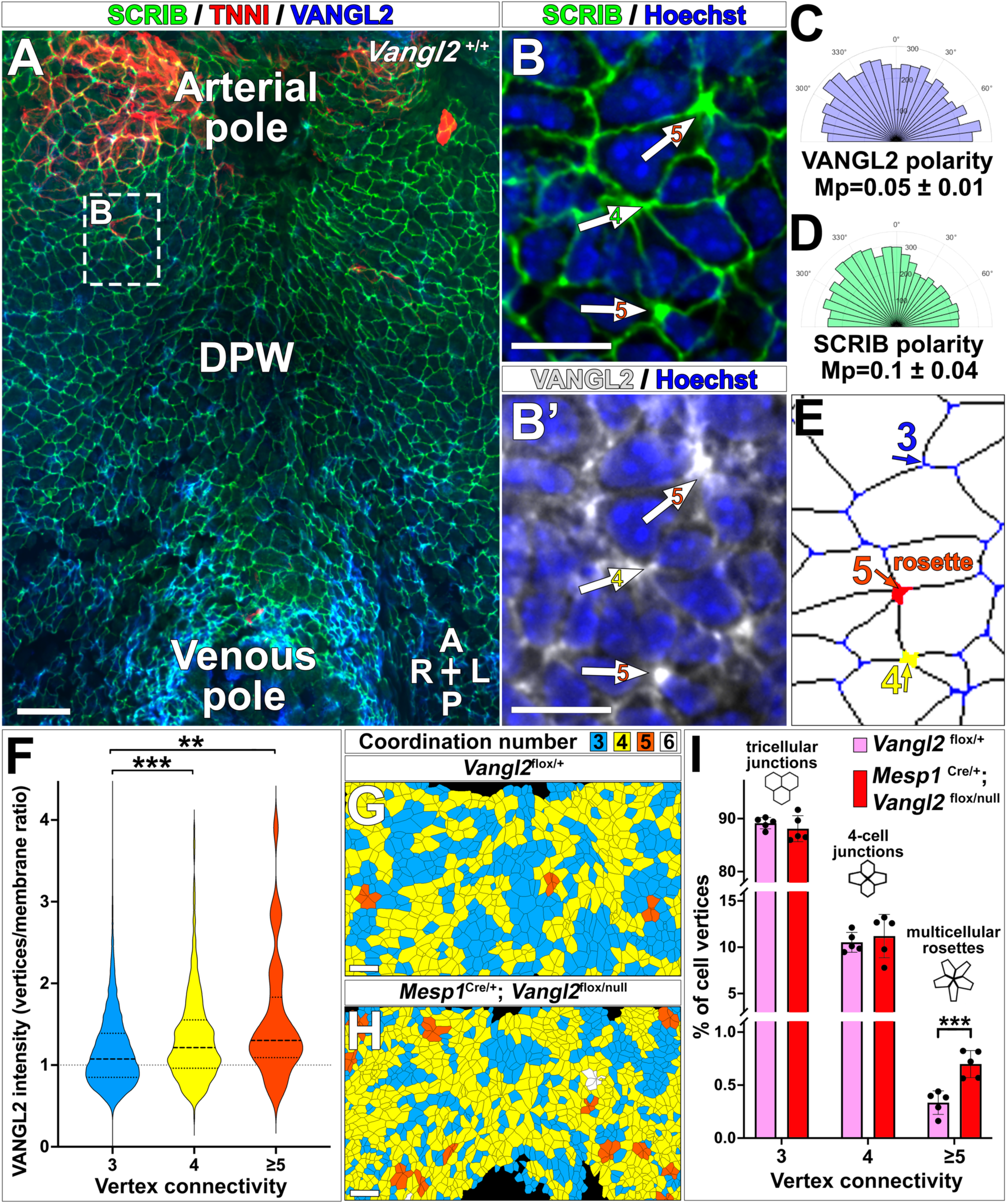
*Vangl2* regulates multicellular rosette resolution rather than planar polarity in the second heart field. (A-B) Cellular localisation of VANGL2 (blue in A, white in B’), compared to the membrane marker SCRIB (green), detected by immunostaining of the micro-dissected dorsal pericardial wall in E9.5 wild-type mouse embryos (n=5) shown in a ventral view. Differentiating cardiomyocytes are marked by TNNI (red). Arrows point to protein accumulation in multicellular junctions, and numbers indicate vertex connectivity. (C-D) Corresponding rose diagrams of VANGL2 and SCRIB planar distribution in cell junctions. The mean polarity value (Mp, n=5 embryos) is provided. (E) Automatic detection of cell vertices and classification according to their connectivity values. Multicellular rosettes correspond to a vertex connectivity value ≥5. (F) VANGL2 intensity in cell vertices was quantified and normalised to cell membrane intensity. *** p value <0.001, **<0.01 (Mann Whitney t-test, n=9848, 730 and 32 vertices with 3, 4 and ≥5 vertex connectivity, respectively, from 5 embryos). (G-H) Analysis of multicellular junctions in the dorsal pericardial wall of controls and *Vangl2* conditional mutants. The coordination number of each cell is colour coded as indicated. (I) Corresponding distribution of multicellular junction classes in *Vangl2* conditional mutants (n=5) compared to littermate controls (n=5). *** p value <0.001 (Fisher’s exact test, n=41 rosettes /12,232 control vertices, n=83 rosettes /11,632 mutant vertices). Means and standard deviations are shown. A, anterior; L, left; P, posterior; R, right. Scale bars : 20 µm (A, F-G), 10 µm (B). See also Figure S4, Source data.

### *Vangl2* is required for multicellular rosette resolution in the second heart field leading to outflow tract elongation

To gain functional insight into the role of *Vangl2* in multicellular junctions, we quantified their complexity in *Vangl2* mutant dorsal pericardial walls compared to littermate controls. We observed a significant increase in multicellular junctions with at least five vertices in *Mesp1*^Cre/+^;*Vangl2*^flox/null^ conditional mutants (Fig. 3G-I). Such accumulation of multicellular rosettes indicates that VANGL2 is required for rosette resolution in the second heart field. We then used the chick embryo to assess whether multicellular rosette resolution could be important for outflow tract elongation. Based on previous observations in the fly and mouse neuroepithelium^42,43^, we selected the *Pten* inhibitor VO-OHpic to impair multicellular rosette resolution. In the chick dorsal pericardial wall, treatment with the *Pten* inhibitor significantly changed the distribution of multicellular junctions with three, four or five vertices, leading to an accumulation of multicellular rosettes indicative of impaired rosette resolution (Fig. 4A-E). The treatment did not significantly affect cell proliferation (Fig. S5A-C). We then tracked progenitor incorporation into the outflow tract by DiI labelling of the dorsal pericardial wall at HH17 (Fig. 4F) followed by 24 hours of *in ovo* development. Compared to control embryos (Fig. 4G-H, Fig. S5D-F), microinjection of the *Pten* inhibitor together with DiI prevented the incorporation of labelled cells into the outflow tract, so that DiI remained in the dorsal pericardial wall (Fig 4I-J, Fig S5G-I). Our results overall show that *Vangl2* is required for outflow tract elongation by promoting multicellular rosette resolution, underlying convergent extension of the second heart field.

**Figure 4.**
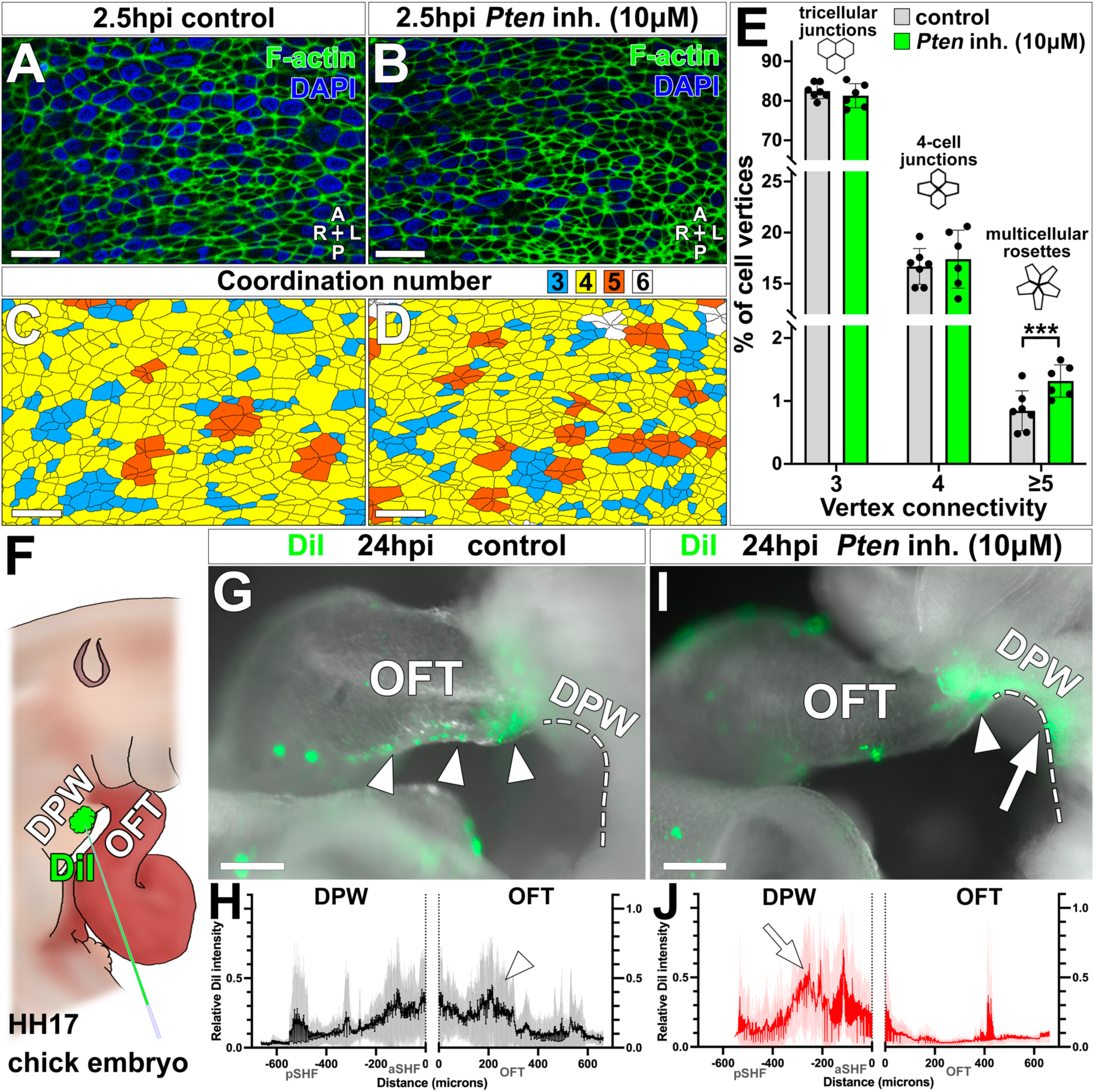
Multicellular rosette resolution in the second heart field is required for outflow tract elongation. (A-B) Staining of micro-dissected dorsal pericardial walls with the cortical marker phalloidin (green) in chick embryos without (A) or with (B) a *Pten* inhibitor (inh.), 2.5h after treatment. (C-D) Corresponding analysis of multicellular junctions, with colour coded cell coordination number. (E) Distribution of multicellular junction classes in control (n=7) and treated (n=6) embryos. *** p value <0.001 (Fisher’s exact test, n=117 rosettes /14,141 control vertices, n=139 rosettes / 10,478 vertices in treated samples). Means and standard deviations are shown. (F) Schematic representation of DiI labelling in the dorsal pericardial wall (DPW). (G-I) Brightfield images of chick embryos in a left lateral view 24h after DiI labelling (green) without (G) or with (I) the inhibitor. Arrowheads and arrow highlight labelling of the inferior outflow tract (OFT) and dorsal pericardial wall, respectively. (H-J) Quantification of relative DiI intensity from the dorsal pericardial wall (negative values) to the outflow tract (positive values), along the dashed lines shown in G (control, n=9) and I (treated, n=5). Accumulation of DiI in the outflow tract (arrowhead) or dorsal pericardial wall (arrow) is indicated. A, anterior; hpi, hours post-injection; L, left; nt, notochord; P, posterior; R, right. Scale bars : 200 µm (J-K), 20 µm (A, F-G, L-O), 10 µm (B). See also Figure S5, Source data.

### *Vangl2* in the *Hoxb1-*positive mesoderm regulates left ventricle position but not outflow tract elongation

Given that *Mesp1*^Cre/+^; *Vangl2*^flox/null^ conditional mutants partially phenocopy *Vangl2^null/null^* constitutive mutants, we generated an alternative conditional mutant with targeted inactivation in more posterior cardiac precursors expressing *Hoxb1*^44,45^. In *Vangl2* constitutive mutants, the position of ventricles is not only affected in terms of orientation along the left-right axis (Fig. 1D-F), but also such that the left ventricle acquires a more right-sided position (Fig. 5A-C). Opposite to *Mesp1*^Cre/+^; *Vangl2*^flox/null^ conditional mutants (Fig. 2C), *Hoxb1*^Cre/+^; *Vangl2*^flox/null^ conditional mutants have normal outflow tract elongation (Fig. 5H), but phenocopy the abnormal position of the left ventricle in constitutive mutants (Fig. 5C, G). The atrioventricular canal is also shifted rightward, indicating an overall rotation of the heart tube around its midline anchor in mutants (Fig. 5A-B, D-E). Thus, *Vangl2* in the posterior *Hoxb1*-positive mesoderm is important to anchor the heart tube and maintain the left ventricle position.

**Figure 5.**
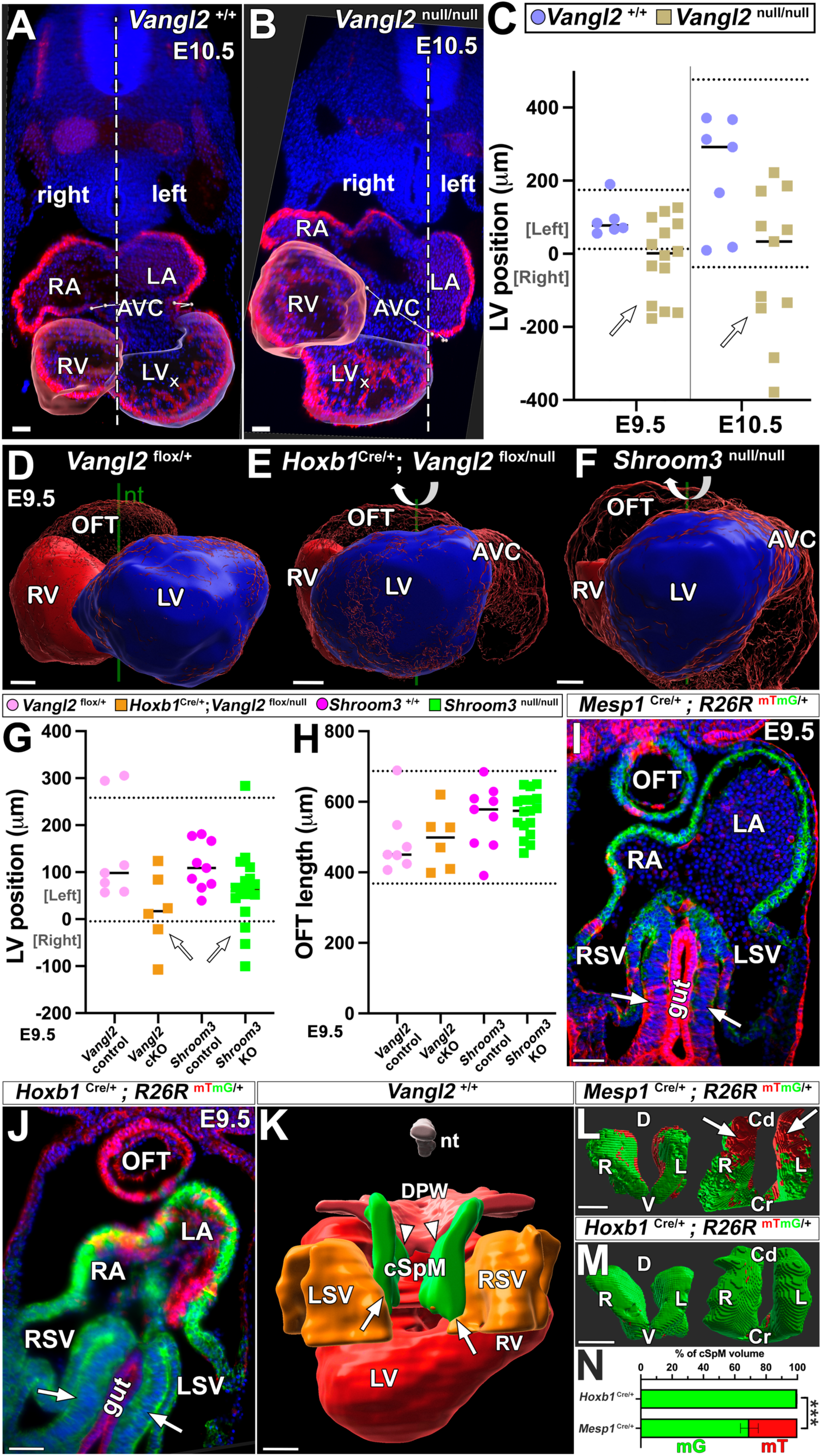
Conditional inactivation of *Vangl2* in the *Hoxb1*-positive mesoderm disrupts left ventricle position, similar to *Shroom3* inactivation. (A-B) Whole mount immunostaining of the myocardium (ACTA2) in E10.5 *Vangl2* constitutive mouse mutants and controls. Segmented ventricles are overlaid on a transversal section. Crosses and dashed lines indicate left ventricle (LV) centroid and embryo midline. (C) Quantification of LV position relative to the midline (n=14 and 11 *Vangl2*^null/null^, 6 and 7 *Vangl2*^+/+^ embryos at E9.5 and E10.5, respectively). (D-F) 3D segmentations of the right (RV, red) and left (blue) ventricles in E9.5 *Vangl2* conditional (E) and *Shroom3* constitutive (F) mutants compared to controls (D). In ventral views, aligned with the notochord (nt), arrows outline the rotated position of mutant hearts. (G) Quantification of LV position (n=7 *Vangl2*^flox/+^, 6 *Hoxb1*^Cre/+^;*Vangl2*^flox/null^, 9 *Shroom3*^+/+^, 17 *Shroom3*^null/null^ embryos). (H) Quantification of outflow tract length (n=7 *Vangl2*^flox/+^, 6 *Hoxb1*^Cre/+^;*Vangl2*^flox/null^, 9 *Shroom3*^+/+^, 17 *Shroom3*^null/null^ embryos). (I-J) Optical frontal sections of E9.5 embryos showing the cells (green) which have expressed *Mesp1* (I) and *Hoxb1* (J), respectively. Arrows point to the splanchnic mesoderm caudal to the heart, which ensheathes the gut. (K) 3D segmentation of the caudal splanchnic mesoderm (cSpM, green) relative to the heart tube (red), sinus venosus (orange) and dorsal pericardial wall (DPW, pink), seen in a caudal view at E9.5. Arrowheads and arrows show its continuity with the dorsal pericardial wall and sinus venosus, respectively. (L-N) The segmented caudal splanchnic mesoderm is labelled by *Hoxb1*^Cre/+^ (M) but partially by *Mesp1*^Cre^ (L, red). (N) Corresponding quantification : 99% and 69%, respectively. *** p value <0.001 (Mann Whitney t-test, n=5 *Mesp1*^Cre/+^*; ROSA26R*^mTmG/+^ and 5 *Hoxb1*^Cre/+^*; ROSA26R*^mTmG/+^ embryos). Means are shown. Dashed lines in C, G-H show the 95% distribution interval based on a Gaussian distribution of control samples. Mutants outside the interval are highlighted by arrows. AVC, atrioventricular canal; Cd, caudal; cKO, conditional knock-out; Cr, cranial; D, dorsal; KO, knock-out; L, left; LA, left atrium; LSV, left sinus venosus; OFT, outflow tract; R, right; RA, right atrium; RSV, right sinus venosus; V, ventral. Scale bars: 100 µm. See also Figure S3, S6, Video S1, Source data.

Given the distinct phenotypes of the two conditional mutants, we explored the intersection between the two Cre drivers. Genetic tracing with the *R26^mTmG/+^* reporter shows that *Mesp1*^Cre/+^ more specifically targets the arterial pole of the heart (e.g. outflow tract) and the entire dorsal pericardial wall (Fig. S3B-D, G). The two drivers overlap in the venous pole of the heart (e.g. atria) (Fig. S3F-I). In contrast, *Hoxb1*^Cre/+^ more specifically targets a region of mesodermal cells caudal and dorsal to the heart tube, continuous with the dorso-medial side of the sinus venosus, and ensheathing the gut, referred to as the caudal splanchnic mesoderm (Fig. 5I-N, Video S1). This indicates that *Vangl2* could play a role caudal to the venous pole of the heart to control the left ventricle position.

### VANGL2 and downstream actin-binding protein SHROOM3 are required for the caudal splanchnic mesoderm architecture associated with left ventricle positioning

We analysed how *Vangl2* regulates the caudal splanchnic mesoderm. We first assessed the cellular localisation of VANGL2 in this tissue and observed an apical enrichment (Fig. 6A). Because the actin-binding protein SHROOM3 acts as a downstream effector of VANGL2 in the mouse neural tube^13^, we assessed its expression using the *Shroom3*^Gt/+^ reporter^19^. *Shroom3* was not detected in the heart tube at E9.5, but it was expressed in the caudal splanchnic mesoderm (Fig. 6B). By immunofluorescence, we found an apical enrichment of SHROOM3 similar to VANGL2 (Fig. 6C). We then analysed the function of *Shroom3* in the embryonic heart. We found that *Shroom3^null/null^* constitutive mutants phenocopy *Hoxb1*^Cre/+^;*Vangl2*^flox/null^ conditional mutants: they display a more right-sided position of the left ventricle, and normal outflow tract elongation (Fig. 5F-H). Our results show that both *Vangl2* and *Shroom3*, which are expressed in the caudal splanchnic mesoderm, control the left ventricle position.

**Figure 6.**
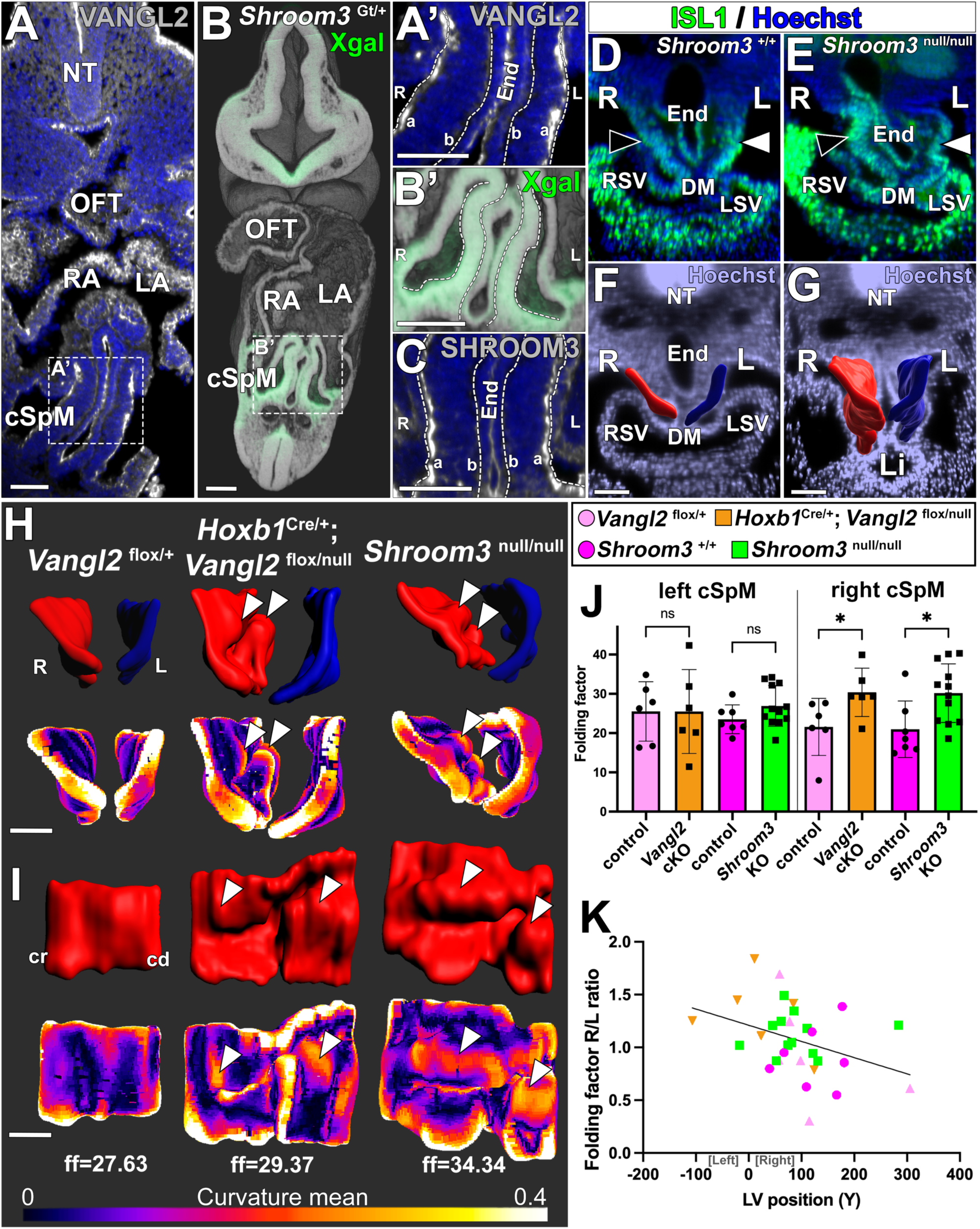
*Vangl2* and *Shroom3* are required to maintain the caudal splanchnic mesoderm architecture, correlating with left ventricle position. (A) Optical transverse section of whole mount VANGL2 immunostaining (white) in a control E9.5 mouse embryo. The caudal splanchnic mesoderm (A’) displays apical (a) enrichment of VANGL2 (n=2). (B) *Shroom3*, detected by Xgal staining (green) in E9.5 *Shroom3*^Gt/+^ embryos imaged by HREM, is expressed in the caudal splanchnic mesoderm (B’, n=7). (C) SHROOM3 immunostaining (white) shows apical enrichment in the caudal splanchnic mesoderm (n=3). (D-E) ISL1 immunostaining in a control (D) and *Shroom3* constitutive mutant (E) at E9.5. The right caudal splanchnic mesoderm (empty arrowheads) is disorganised in *Shroom3* constitutive mutants. (F-G) 3D segmentation of the right (red) and left (blue) caudal splanchnic mesoderm, from the dorsal mesocardium (DM) to the liver primordium (Li), in a E9.5 control. Cranial (F) and caudal (G) transverse sections are shown. (H-I) Segmentation of the caudal splanchnic mesoderm (right, red ; left, blue) in *Vangl2* conditional and *Shroom3* constitutive mutants compared to controls, shown in transverse (H) and midline (I) views. Arrowheads point to increased folding on mutant right sides. The mean tissue curvature is colour coded below. The folding factor (ff) representing the ratio between high and low curvature is provided. (J) Distribution of the folding factor in the right and left caudal splanchnic mesoderm (cSpM). * p value < 0.05 (Mann Whitney t-test, n=6 *Vangl2*^flox/+^, 6 *Hoxb1*^Cre/+^;*Vangl2*^flox/null^, 7 *Shroom3*^+/+^, 12 *Shroom3*^null/null^ embryos). (K) Correlation between the folding factor asymmetry in the caudal splanchnic mesoderm and the position of the left ventricle (LV) in *Vangl2* conditional (orange downward triangles), *Shroom3* constitutive (green squares) mutants and controls. * p value < 0.05 (Spearman correlation, n=31). Means and standard deviations are shown. a, apical side; b, basal side; cd, caudal side; cKO, conditional knock-out; cr, cranial side; End, endoderm; KO, knock-out ; L, left; LA, left atrium; LSV, left sinus venosus; NT, neural tube; ns, non-significant; OFT, outflow tract; R, right; RA, right atrium; RSV, right sinus venosus;. Scale bars: 100 µm. See also Figure S6-S7, Source data.

We then investigated whether the caudal splanchnic mesoderm was affected in mutants. In controls, the caudal splanchnic mesoderm is a regular epithelium (Fig. 6D), whereas it appeared disorganised and folded in mutants (Fig 6E). We segmented (Fig. 6F-G) and quantified its shape and found significantly increased folding on the right side, both in *Shroom3^null/null^* constitutive mutants and *Hoxb1*^Cre/+^;*Vangl2*^flox/null^ conditional mutants (Fig 6H-J). This was not the case in *Mesp1*^Cre/+^;*Vangl2*^flox/null^ conditional mutants, which have a normal caudal splanchnic mesoderm and a normal position of the left ventricle (Fig S6). The right over left ratio of the folding factor in the caudal splanchnic mesoderm significantly correlates with the lateral position of the left ventricle relative to the midline (Fig 6K). In keeping with the asymmetric disorganisation of the caudal splanchnic mesoderm architecture, we observed a slight bias in *Shroom3* expression, with right-sided enrichment (Fig. S7). Our observations thus show that *Vangl2* and *Shroom3* are important to maintain the architecture of the right caudal splanchnic mesoderm, which is associated with the position of the left ventricle.

## Discussion

Using a series of mutants and quantitative phenotyping, we demonstrate that *Vangl2* plays a dual role to shape the mouse embryonic heart tube (Fig. 7). By promoting multicellular rosette resolution in the dorsal pericardial wall, it drives cell rearrangements underlying convergent extension of the second heart field, leading to elongation of the tube arterial pole. At the other, venous, pole, it rather maintains the caudal splanchnic mesoderm architecture together with its downstream effector SHROOM3, and thus can indirectly prevent rotation of the tube around its midline anchor. We thus uncover a process of heart morphogenesis, whereby bilateral symmetry of a tissue outside the heart needs to be maintained for correct ventricle position. Whereas PCP has been mainly analysed in planar tissues or 2D tissue sections, we provide an example of how VANGL2 can shape an organ in 3D.

**Figure 7.**
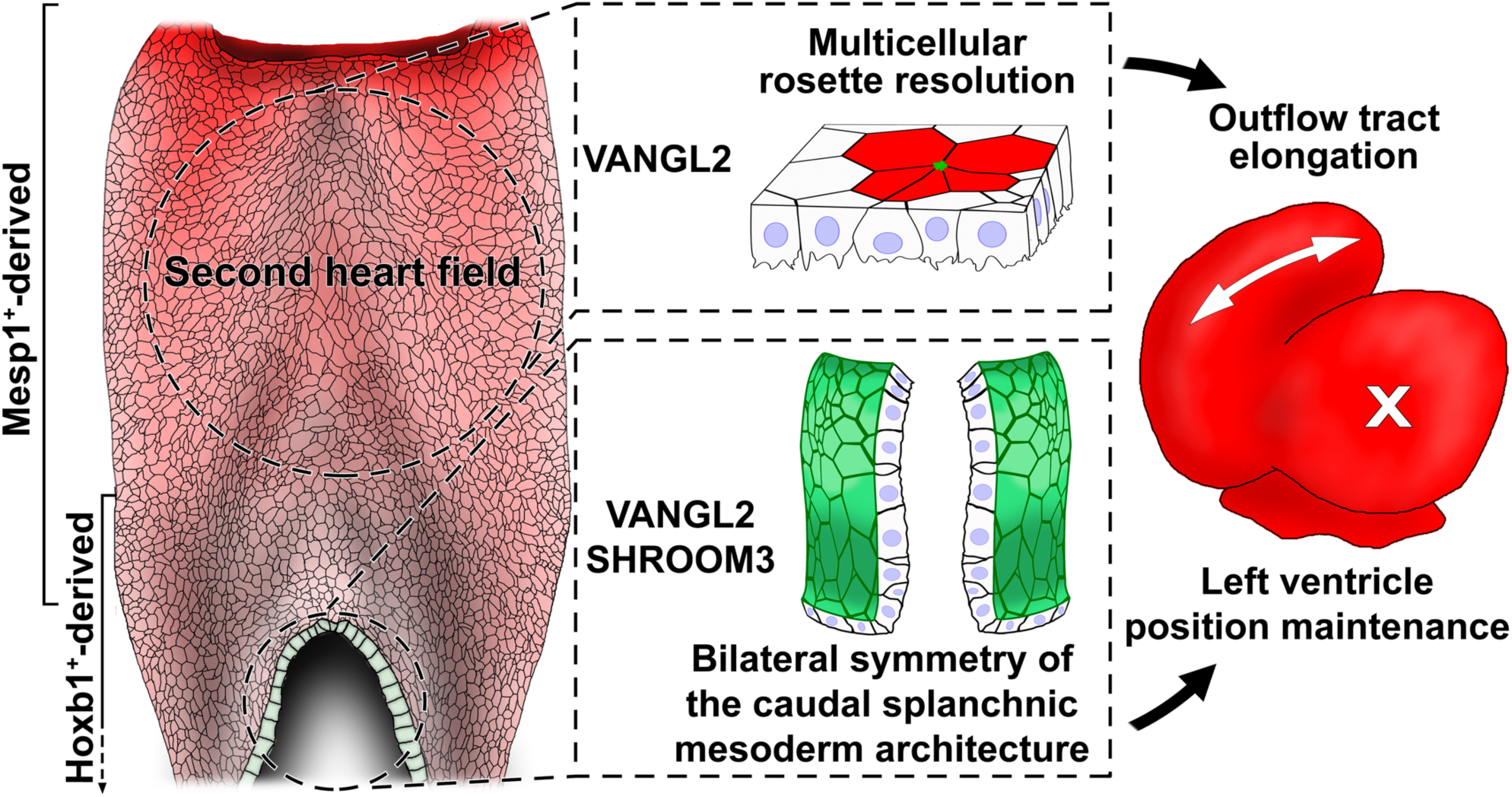
Model of *Vangl2* role in shaping the embryonic heart tube. In the second heart field epithelium, located in the dorsal pericardial wall, VANGL2 is not planar polarised, but rather accumulates in multicellular junctions. It is important for rosette resolution, thus promoting cell rearrangements involved in the elongation of the outflow tract, i.e. the arterial pole of the heart tube. In another epithelium, the caudal splanchnic mesoderm, to which the venous pole of the heart is attached, VANGL2 is enriched at the apical side, together with its downstream actin binding effector SHROOM3. Inactivation of *Vangl2* or *Shroom3* disrupts the architecture of the right caudal splanchnic mesoderm. Loss of bilateral symmetry results in a rotation of the heart tube around the midline, shifting the left ventricle to the right. The *Mesp1^Cre^* and *Hoxb1^Cre^* drivers target the second heart field epithelium and caudal splanchnic mesoderm, respectively. Our model shows how *Vangl2* plays a dual role, unrelated to planar polarity, in shaping the developing heart tube from its two poles.

*Vangl2* is important to shape the mouse heart tube. In the constitutive mutant, because the ventricles have not reached their definitive left-right position, it had been proposed that *Vangl2* controls heart looping^17^. However, we do not find defects in the core mechanism of heart looping, i.e. buckling between E8.5 and E9.5^24^. In addition, heart defects are more pronounced at E10.5, after heart looping is completed. We further analyse previous qualitative observations of outflow tract elongation defects^20^. Using quantitative approaches and 3D imaging, we have assessed the morphological and cellular phenotypes of *Vangl2* mutants with higher resolution. VANGL2 subcellular localisation is now analysed in the plane of the second heart field, compared to the apico-basal axis previously. We thus uncover that VANGL2 does not show any planar polarisation but is involved in planar cell rearrangements by controlling multicellular rosettes. In another mutant, for *Greb1l*, we had shown how the maintenance of a cardiac precursor pool in the second heart field is required for outflow tract elongation^27^. Yet, *Greb1l* mutants display a more severe cardiac phenotype, with 40% decrease in outflow tract length at E10.5, compared to 13% in *Vangl2* mutants. This reflects distinct cellular roles: *Greb1l* is required for ribosome biogenesis and cell differentiation, whereas *Vangl2* for cell rearrangements. Convergent extension of the second heart field had been proposed based on cell tracing experiments^34,35^. We now provide a cellular mechanism, multicellular rosettes, and a regulator, VANGL2. Another member of the PCP pathway, WNT5A, was also shown to be required for second heart field convergent extension^36^. However, the genetic interaction between *Vangl2* and *Wnt5a* in this process remains to be demonstrated. The role of *Vangl2* in the second heart field is consistent with its regulation of convergent extension in other tissues like the mouse body axis, mouse neural plate and worm ventral nerve cord^11,15,16^.

*Vangl2* is required in different tissues for heart development. Extending previous work^20^, we use alternative Cre lines to further dissect the cell populations that require *Vangl2* for shaping the heart tube. We uncover the caudal splanchnic mesoderm, labelled by *Hoxb1^Cre^*, as an epithelium anchoring the heart tube. This tissue, which is continuous with the second heart field and sinus venosus, has been previously related to the heart because it contains cardiopulmonary progenitors^29^. It surrounds the endoderm and gives rise to the smooth muscle and cartilage of the airways^46^. We now show that it can indirectly modulate ventricle position, by acting as a midline anchor. This reveals an additional aspect of heart morphogenesis. The observation that *Shroom3* mutants phenocopy *Vangl2* conditional mutants is reminiscent of their genetic interaction during closure of the neural tube^13^. There, *Shroom3* regulates epithelial cell shape by controlling the actomyosin network associated with the apical junctional complex^12^. This suggests a potential mechanism regulated by *Vangl2* and *Shroom3* to maintain the cellular architecture of the caudal splanchnic mesoderm.

VANGL2 function differs between tissues. *Van Gogh* was initially identified as an important regulator of planar cell polarity in the fly^47^. This role is conserved in mammalian epithelia such as the epidermis, cochlea, multiciliated tracheal and ependymal cells^6,48–50^. Planar polarity of VANGL can manifest in mesenchymal tissues, as shown in the limb^51^. We detect an apical localisation of VANGL2 in the caudal splanchnic mesoderm, which is in keeping with the involvement of VANGL not only in planar, but also apico-basal polarity, i.e. in the perpendicular axis^9,52–54^. Similar to our finding in the second heart field, VANGL can function outside the core PCP complex in the lung mesenchyme. During airway branching VANGL1/2 are required for the cellular architecture of the airway smooth muscle^10^, reminiscent of our observations in the caudal splanchnic mesoderm. At later stages of lung development, VANGL1/2 is required for mesenchymal cell shape and tissue fluidity during sacculation^40^. In the worm, VANG-1, which is required for rosette resolution, is localised at multicellular junctions and acts together with Robo^16^, suggesting a potential mechanism for VANGL2 mediated rosette resolution in the second heart field. Thus, VANGL can play several roles, related or not to polarity, to regulate cell behaviour such as orientation, shape, motility, intercalation, based on cytoskeleton and junction remodelling^5^.

The phenotype of *Vangl2* mutants shows a striking left-right asymmetry in the caudal splanchnic mesoderm. *Vangl1* and *Vangl2* have previously been involved in left-right asymmetry, but at the early stage of symmetry-breaking in the node^55^. They regulate the planar polarisation of the basal body, which is important for the coordinated tilting of motile cilia and thus the generation of a directional fluid flow. During organogenesis, the PCP pathway was shown to be important for gut looping in the fly and chick, by regulating asymmetric cell behaviour^56,57^. However, in the caudal splanchnic mesoderm, *Vangl2* is not required for asymmetric cell behaviour, but rather to maintain symmetry. This is reminiscent of the buffering mechanism identified in the presomitic mesoderm, whereby retinoic acid signalling counteracts the asymmetric expression of the left determinant *Nodal*, to maintain somite bilateral symmetry^58,59^. Whether *Vangl2* and *Shroom3* also interact with *Nodal* remains to be investigated. In the context of the heart, *Nodal* is known to be important for controlling asymmetric cell behaviour and biasing the buckling mechanism of heart looping^25^. The importance of bilateral symmetry in the caudal splanchnic mesoderm provides a novel regulatory process during asymmetric heart morphogenesis. This is potentially relevant to patients, in which *SHROOM3* variants are associated with the heterotaxy syndrome, corresponding to left-right anomalies in visceral organs^60^. Overall, our dissection of the developmental mechanisms regulated by VANGL2 and its partner provides novel insight into the origin of congenital heart defects.

## Acknowledgements

We thank L. Guillemot, V. Benhamo, L. Borges, J. Terret, N. Agueeff, M. Meunier, E. González-Muñoz, A. Sánchez-Mata for technical assistance; R. Levayer for insightful discussions ; T. Holm Bønnelykke, E. Perthame, S. Paramore, L.A. Davidson, B. Aigouy for expert advice; D.J. Henderson for providing *Vangl2^flox/+^* mice, T. Drysdale for providing *Shroom3^flox/flox^* mice; M. Durbin for providing *Shroom3^Gt(ROSA^*^53^*^)Sor^* embryos, T. Nishimura and M. Takeichi for providing SHROOM3 antibody; D. Conrozet and the histology platform of the SFR Necker; S. Dupichaud, M. Garfa-Traore and the Cell imaging platform; N. Goudin of the Necker Bio-image analysis plateform; the LEAT animal facility. This work was supported by core funding from the Institut Pasteur and INSERM, state fundings from the Agence Nationale de la Recherche under ‘‘Investissements d’avenir’’ program (ANR-10-IAHU-01, ANR-10-LABX-73-01 REVIVE), an Equipe FRM grant, the Philanthropy Department of Mutuelles AXA through the Head and Heart Chair to S.M.M. and the Spanish Ministry of Science-AEI grant PID2021-122626OB to JMPP. A.U.H. was supported by the Fulbright US France programme. G. L. is a CNRS Research Engineer. P.P-G. was supported by fellowships from Pasteur Roux-Cantarini, and the Fondation Lefoulon-Delalande. S.M.M. is an INSERM research director.

## Author contributions

Conceptualization, P.P.G. and S.M.M.; Formal analysis, P.P.G., G.L.; Funding acquisition, P.P.G., S.M.M. and J.M.P.P.; Investigation, P.P.G. and A.U.H; Methodology, P.P.G., G.L.; Software, G.L.; Resources, J.M.P.P.; Supervision, P.P.G. and S.M.M.; Visualization, P.P.G.; Writing – original draft, P.P.G. and S.M.M.; Writing – review & editing, all authors.

## Methods

### EXPERIMENTAL MODEL

#### Animals

*Vangl2^null/+^* mice were generated by crossing *Vangl2^tm1.1Djhe/tm1.1Djhe^* (referred to as *Vangl2^flox/flox^* ^20^) males with *Tg Mef2cAHFCre* transgenic females^61^. *Shroom3^null/+^* mice were generated by crossing *Shroom3^flox/flox^* males^62^ with *Tg Mef2cAHFCre* females. *Vangl2*^flox/flox^ and *Vangl2*^null/+^ lines were maintained in a C57Bl/6J genetic background, while *Shroom3*^null/+^ line was maintained in a C57Bl/6N background. *Vangl2*^null/+^ females were crossed with *Mesp1*^Cre/+ 38^ and *Hoxb1*^Cre/+ 63^ males to obtain *Mesp1*^Cre/+^ ; *Vangl2*^null/+^ and *Hoxb1*^Cre/+^ ; *Vangl2*^null/+^ studs, respectively. These studs were crossed with *Vangl2*^flox/flox^ females to obtain *Mesp1*^Cre/+^;*Vangl2*^flox/null^ and *Hoxb1*^Cre/+^;*Vangl2*^flox/null^ conditional mutant embryos, respectively. *Gt(ROSA)26Sortm4(ACTB-tdTomato,-EGFP)Luo* (referred to as *R26^mTmG^*, ^64^) were used to control Cre expression profiles. *Shroom3^Gt(ROSA53)Sor^* are referred to as *Shroom3*^Gt/+ 14^.

Both male and female embryos were collected and used blindly for experiments. Control embryos are wild-type littermates. Embryonic day (E) 0.5 was defined as noon on the day of vaginal plug detection. All embryos were genotyped by PCR of the yolk sac, using primers listed in Table S1. For the genotyping of *Shroom3* alleles^62^, primers were designed to detect the wild-type (282bp), floxed (316bp) and deleted (217bp) alleles. Animals were housed in the Laboratory of Animal Experimentation and Transgenesis of the SFR Necker, Imagine Campus, Paris. Animal procedures were approved by the ethical committee of Université Paris Cité and the French Ministry of Research.

Fertilized chicken (*Gallus gallus*) eggs were kept in an incubator at 38°C. Embryos were staged according to the Hamburger and Hamilton stages of chick development^65^. Animals were handled in compliance with the international guidelines (1964 Declaration of Helsinki) for animal care and welfare. Experimental procedures have been revised and approved by the Ethics Committee at the University of Malaga.

## METHOD DETAILS

### In ovo injection of chick embryos

VO-Ohpic trihydrate (Selleckchem) was used as a potent inhibitor of Pten, stored at 25mM in DMSO. It was injected *in ovo* after dilution in PBS at 10µM, and mixing with 200 µg/ml DiI celltracker. The inhibitor and adjuvant solutions were injected on the dorsal pericardial wall of chick embryos at HH17 stage (2.5 days of incubation) using an Eppendorf FemtoJet 4i Microinjector. Eggs were closed for further incubation of either 2.5 hours or 24 hours. Embryos were then micro-dissected in PBS and fixed in 4% PFA overnight at 4°C. Fluorescent images of embryos were acquired under a Leica stereoscope. The dorsal pericardial wall of HH17 chick embryos 2.5 hours post injection was then micro-dissected for immunostaining.

#### Whole mount immunostaining

For ISL1/ACTA2 immunostaining, whole E9.5 mouse embryos were incubated in cold 250mM KCl cardioplegia solution for 1 minute and fixed in 4% cold PFA overnight. Samples were washed in PBS, gradually dehydrated in methanol (25%, 50%, 75% in PBS and 100%) and stored at -20°C. Samples were then rehydrated and washed in PBS at room temperature. Embryos were cleared using an adapted CUBIC protocol^24^. Shortly, samples were incubated in a 1:1 solution of water:CUBIC reagent-1 for 1 hour at 37°C and in pure reagent-1 overnight at 37°C. Then samples were washed in PBS and incubated with primary antibodies (2-5 µg/ml). ISL1 (DSHB) and ACTA2 (Abcam) primary antibodies were diluted in TNB blocking solution with 0.5% triton for 48-76h at 4°C. Samples were then washed in PBS (4x30min at room temperature) and incubated with secondary antibodies (1/500) and Hoechst (1/2000) for 24- 48h at 4°C. Samples were then gently washed in PBS and incubated in CUBIC Reagent-2 diluted in PBS (1:1) for 4 hours at 37°C and then in Reagent-2 overnight at 37°C. Cleared embryos were then mounted in warm R2:agarose solution in separated capillaries. Multi- channel 16-bit images were acquired with a Z.1 lightsheet microscope with a 20X/1.0 objective.

For SHROOM3 and VANGL2 immunostaining, whole E9.5 mouse embryos were fixed in 4% PFA for 45 minutes on ice. VANGL2 (1/200, Millipore) and SHROOM3 (1/500 ^66^) were diluted in TNB blocking solution with 0.5% triton and 1% fish gelatin. The TSA amplification kit was used to enhance signal intensity. Samples were bleached by a 30 minute incubation in 3% H_2_O_2_. They were washed in PBS and incubated in biotin-conjugated secondary antibodies (1/500) overnight at 4°C. Samples were treated with streptavidin-HRP overnight at 4°C and tyramide-Cy3 7 minutes at room temperature. SHROOM3 immunostaining is not compatible with the traditional CUBIC reagent-1, so we used the alternative Reagent-1A^67^. Embryos were cleared and imaged as above.

E9.5 mouse neural tubes or dorsal pericardial walls of embryos were microdissected^68^ and fixed in 4% PFA for 45 minutes on ice^6^. Samples were washed in TNB with 0.5% triton and 1% fish gelatin for 30 minutes at room temperature. Primary antibodies against VANGL2, SCRIB (Santa Cruz) and TNNI (Santa Cruz) were diluted at 2-5 µg/ml in TNB with 0.5% triton and 1% fish gelatin were added for 48-76h at 4°C. For VANGL2 staining, samples were bleached in 3% H_2_O_2_ for 30 minutes, washed in PBS, incubated with a biotin-conjugated secondary antibody (1/500) overnight at 4°C, washed and incubated with Tyramide Cy3 TSA amplification kit for 7 minutes. For SCRIBBLE and TNNI staining, samples were incubated with fluorescent secondary antibodies (1/500) overnight at 4°C. Multi-channel 16-bit images were acquired with a confocal spinning disk microscope.

Control and treated dorsal pericardial walls of chick embryos were washed in PBS and incubated in SBT blocking solution (16% sheep serum, 1% bovine serum albumin and 0.5% TritonX-100 in PBS) for 1 hour. Samples were then incubated overnight at 4°C with a primary antibody against PHH3 (1/200), then 2 hours at room temperature with a FITC conjugated secondary antibody (1/500), DAPI (1/2000) and AF647-conjugated phalloidin (1/400). Samples were washed, mounted in 75% glycerol and imaged under a SP5 laser confocal microscope.

#### Whole mount endogenous fluorescence

E9.5 *R26^mTmG^* embryos crossed with a Cre line were selected for their endogenous green and red fluorescence and fixed in 4% PFA for 1 hour at room temperature. Embryos were washed in PBS and cleared using the CUBIC approach. Samples were incubated in a 1:1 solution of water:CUBIC reagent-1 for 1 hour at 37°C and in pure reagent-1 overnight at 37°C. They were washed in PBS, incubated with Hoechst (1/2000) for 2 hours and washed again in PBS. Samples were then incubated in a 1:1 solution of CUBIC Reagent-2 and PBS for 4 hours at 37°C, and in Reagent-2 overnight at 37°C. Samples were embedded in a solution of 10% low melting agarose diluted in reagent2 and mounted in capillaries. The refractory index of the R2-agarose solution was controlled to match that of reagent 2 (1.45). Multi-channel 16-bit images were acquired with a Z.1 Lightsheet microscope.

#### X-gal staining

*Shroom3*^Gt/+^ E9.5 embryos were genotyped and stained with X-gal as previously^19^. *Shroom3*^Gt/+^ embryos were then embedded in methacrylate resin (JB4) containing eosin and acridine orange as contrast agents. Two channel images of the surface of the resin block were acquired using an optical high-resolution episcopic microscope with a 1X Apo objective, repeatedly after removal of 1.7 μm thick sections. The tissue architecture was imaged with a GFP filter and the X-gal staining with a RFP filter. The 3D dataset comprises 1100-1300 images of 1.5-1.7 μm resolution in x and y. Fiji (ImageJ) software was used to crop or scale the datasets. Tiff files were transformed in Imaris files using Imaris File Converter tool.

## QUANTIFICATION AND STATISTICAL ANALYSIS

### Quantifications of the geometry of the heart tube

Heart tube segmentation and geometry quantification in whole mount immunostained embryos was adapted from previous work^27^. In 3D lightsheet images, the myocardium contour of four heart tube regions was manually segmented using the IMARIS software: distal outflow tract, proximal outflow tract, right ventricle and left ventricle. The boundary between the outflow tract and right ventricle is based at the transition between endocardial cushions and myocardium trabeculation. The outflow tract length was measured in 3D along the line joining five successive centroids. Three centroids were obtained with the IMARIS oblique slicer function from three different polygons (MATLAB geom3d library: function polygonCentroid3d) perpendicular to the tube: one at the exit of the outflow tract, one at the junction between the distal and proximal outflow tract segments and one at the boundary between the outflow tract and the right ventricle. Two additional landmarks were obtained by calculating the centroid of the dOFT and pOFT 3D segmentations using IMARIS. The full length of the outflow tract is computed as the cumulative distance along the five centroids.

Samples were aligned in 3D using two landmarks on the notochord and two landmarks in the neural tube, defining the Z and X reference axes, respectively. We then used an in-house MATLAB code^27^ to adjust x and z-axes at 90° angle, align the notochord with the Z-axis and the neural tube with the X-axis and to rotate all segmented landmarks in the heart tube accordingly. The orientation of the right ventricle - left ventricle axis was calculated based on the vector joining the centroids of each ventricle, projected on the transversal plane (XY), and its angle with the notochord.

#### Quantification of tissue proportion targeted by Cre

The second heart field in the dorsal pericardial wall and myocardial regions of the heart tube were manually segmented using the IMARIS software. The outflow tract and caudal remnant of the dorsal mesocardium were used as cranial and caudal landmarks of the dorsal pericardial wall, respectively. The contour of the dorsal pericardial wall and myocardium were reconstructed in 3D by the “Create Surface” function. Signal of the green and red channels intersecting with the different segmented regions were obtained using the ‘‘mask selection’’ function to extract new channels and create new 3D objects (one red and one green for each segmented region). The volume of each was extracted and the proportion of green over green+red was calculated for each region of interest.

#### Segmentation and quantification of the cell architecture

To improve visualization and cell segmentation of the curved dorsal pericardial wall, a surface of interest was generated using the subapical SCRIB signal and was projected using the Tissue Cartography method^69^, both in the SCRIB and VANGL2 channels. Visualization of TNNI and Hoechst staining corresponds to a max intensity projection. Cells were automatically segmented using the CellPose tool and mistakes in the segmentation were manually corrected using the Tissue Analyzer tool^70^ in FIJI.

The coordination number of a cell was used to count and visualize multicellular rosettes, taken as the maximum number of cell neighbours associated with its vertices^13^. A vertex is a point at which 3 or more cells join together. We calculated the connectivity of each vertex as the number of cells joining at this vertex. As the resolution of a vertex is not exactly one pixel (due to imaging resolution and segmentation precision), we considered cells to be joining at a given vertex if they were within a very small radius of 3 pixels around it. We implemented a Fiji^71^ macro Rosette-finder to extract these parameters directly from the “vertices.tif” and “cell_identity.tif” files of Tissue Analyzer. For each segmented cell, we assigned its coordination number as the maximum connectivity of its associated vertex. We calculated the percentage of tricellular junctions, four cell junctions and multicellular rosettes corresponding to a vertex connectivity value of 3, 4 and ≥5, respectively.

A second macro Vertices-intensities was developed to sort cell vertices by their connectivity value and quantify their relative VANGL2 staining. Each vertex (one pixel) was extended to a circle of 2-pixel radius to measure the mean staining intensity in it. All values of the relative VANGL2 intensity for each vertex were represented in a violin plot graph using the Graphpad Prism software.

We quantified the mean polarity value of VANGL2 in cell membranes using the Tissue Analyzer tool^39,70^. The Matlab polar histogram function was used to plot rose diagrams.

To assess cell proliferation, we manually counted with the FIJI multi point tool all nuclei and PHH3-positive nuclei in a 2D image of the flat chick dorsal pericardial wall and calculated the percentage of PHH3-positive nuclei.

#### Quantification of DiI-positive cell ingression into the heart tube

We quantified the DiI intensity along the dorsal pericardial wall and outflow tract using the “plot profile” tool in FIJI. Each intensity measurement was normalized to the maximum intensity value measured for each embryo (pericardial wall and outflow tract). Data were plotted using the Prism Graphpad software.

#### Quantification of the curvature of the caudal splanchnic mesoderm

We segmented the lateral splanchnic mesoderm from the dorsal mesocardium to the liver bud using Imaris. We then measured the mean curvature (Cmean) as previously reported^72^. Based on the mean curvature values, we computed a folding factor (ff), taken as the ratio of the Cmean for the highly curved surface (0.125-0.3), to the Cmean for the least curved surface (0- 0.05).

#### Quantifications of *Shroom3* expression in the caudal splanchnic mesoderm

We segmented the caudal splanchnic mesoderm of *Shroom3*^Gt/+^ embryos and quantified the overall Xgal intensity in 3D. We used the Imaris orthogonal slicer to visualize the most caudal region of the tissue, at the level of the liver bud, and quantified the intensity along the right and left caudal splanchnic mesoderm with the “plot profile” tool in FIJI.

#### Bioinformatic analyses of published single cell RNA sequences

The single cell expression and annotation data (cluster and meta-cell identifiers) of mouse E9.25 microdissected cardiac region were downloaded from GEO (GSE205950) according to https://github.com/gonzalezdavidmatthew/Dubois_embryo_cardiac_seq. The expression profile of reference genes was assessed to annotate each cluster as in the original publication. *Rgs5, Crabp1, Flna* and *3632451O06Rik* expression define cluster 17 as outflow tract; *Isl1* and *Pdgfra* define clusters 2 and 18 as second heart field; *Tbx1* defines cluster 18 as the anterior second heart field, while *Aldh1a2*, *Hoxb1* and *Foxf1* define cluster 2 as the posterior second heart field. More markers of the posterior second heart field, *Tbx5*, *Osr1* and *Foxf1* similarly labeled cluster 2. Gene expression in clusters 17 (OFT; 505 cells), 18 (aSHF; 455 cells) and 2 (pSHF; 1789 cells) was represented in violin plots.

#### Statistical Analyses

Statistical tests, p-values and data points are described in figure legends and Source data. P-values less than 0.05 were considered statistically significant. The collection of full litters was used to randomise imaging experiments. Group allocation was based on PCR genotyping. Exact sample numbers (n) are indicated in the text and refer to biological replicates, i.e. different embryos or different cells. Investigators were blinded to allocation during imaging, but not during quantifications. Too young E9.5 embryos, with less than 18 somites, were discarded from the analysis. Tests were performed with GraphPad Prism and R. Fisher’s exact test was used to compare proportions. Mann-Whitney test was used to compare the mean between two groups, and the one sample t-test to compare it with an expected value. Correlation between two data series was tested using Spearman correlation. The regression line was computed using the least square method. The 95% distribution interval for the control mean was calculated after validation of a normal distribution.

**Correspondence** and requests for materials should be addressed to Sigolène Meilhac.

## Data availability

Data supporting the findings of this study are available in the article, its Supplementary information and the Source Data file. Images generated in this study and used for quantifications have been deposited in the Owey platform (doi: 10.48802/owey._DJu1Zfj.1.0) and will be publicly available as of the date of publication.

## Code availability

Original codes are publicly available at Zenodo (doi : 10.5281/zenodo.8270279) and gitlab (https://gitlab.pasteur.fr/gletort/rosetteAnalysis/).

